## Supplemental Table 1 for "Population dynamics of the primary malaria vector *Nyssorhynchus darlingi* in a high transmission setting dominated by fish farming in western Amazonian Brazil"

**S1 Table.** **Independent variable and response options (by classification or instrument measure), level of subject analysis (sampling-point or fishpond), and sampling effort during 2017 according to the sampling schedule (6-month for Feb, Mar, Apr, May, Aug, Sept; 3-month for Feb, Mar, Apr; 3-month for May, Aug, Sept).**

| **Independent**  **Variable** | **Response** | **Level of subject analysis** | **Sampling Effort** |
| --- | --- | --- | --- |
| Periodicity | permanent or temporary | fishpond | Feb-Sept (6 months) |
| Abandoned | yes or no | fishpond | Feb-Sept (6 months) |
| Associated Vegetion | if present: emerging, submerged, floating | sampling-point | Feb-Sept (6 months) |
| Fauna Presence | *Culex* sp.; amphibians; fish | sampling-point | Feb-Sept (6 months) |
| Physical-Chemistry  (continuos values) | pH; temperature (ºC); conductivity (µS/cm) | sampling-point | Feb-Apr (3-month) |
| Turbity | 0 - 200 (JTU)  (where 0 represents translucent water) | fishpond | May-Sept (3-month) |
| Shadding | 0 – 24.96 (1⁄4"-squares)  (where 0 represents shaded and 24.96 represents completely exposed) | sampling-point | May-Sept (3-month) |
| Physical-Chemistry  (categorical values) | pH, nitrates (mg/L); nitrites (mg/L); carbonate hardness (KH); dissolved chlorine (mg/L) | fishpond | May-Sept (3-month) |
