## Supplemental Table 2 for "Population dynamics of the primary malaria vector *Nyssorhynchus darlingi* in a high transmission setting dominated by fish farming in western Amazonian Brazil"

**S2 Table. Fishpond numbers by site and collection month in Mancio Lima, Acre, Brazil 2017.**

| **Collection**  **Month** | **Site** | | | | | | | | |
| --- | --- | --- | --- | --- | --- | --- | --- | --- | --- |
|  | **1** | | | **2** | | | **3** | | |
|  | **fishpond** | **dry** | **not surveyed** | **fishpond** | **dry** | **not surveyed** | **fishpond** | **dry** | **not surveyed** |
| February | 15 (15) | 0 | 0 | 22 (22 ) | 0 | 0 | 9 (9 ) | 0 | 0 |
| March | 11 (26 ) | 0 | 0 | 4 (26 ) | 0 | 0 | 1 (10 ) | 0 | 0 |
| April | 1 (26 ) | 1 | 0 | 0 (25 ) | 1 | 0 | 0 (9 ) | 0 | 1 |
| May | 0 (25 ) | 2 | 0 | 0 (26 ) | 0 | 0 | 0 (9 ) | 0 | 1 |
| August | 0 (23 ) | 3 | 1 | 0 (22 ) | 4 | 0 | 0 (10 ) | 0 | 0 |
| September | 0 (24 ) | 3 | 0 | 0 (22 ) | 4 | 0 | 0 (8 ) | 2 | 0 |

Fishpond column: numbers of fishponds identified per respective month; values in brackets are total fishponds surveyed for the respective month. Dry column: fishponds with absence of water (by seasonality or owner management). Not surveyed column: fishponds unsampled (due to flooded fields or no possible access to property).
