## Supplemental Table 3 for "Population dynamics of the primary malaria vector *Nyssorhynchus darlingi* in a high transmission setting dominated by fish farming in western Amazonian Brazil"

**S3 Table. Environment independent variables by site in Mancio Lima, Acre, Brazil 2017.**

| **Study Site**  **(n total)** | **Periodicity*** | | **Abandoned** | | **Associated Vegetation**** | | | **Presence**** | | |
| --- | --- | --- | --- | --- | --- | --- | --- | --- | --- | --- |
|  | **Temporary** | **Permanent** | **Yes** | **No** | **Emerging** | **Submerged** | **Floating** | ***Culex* sp.** | **Amphibian** | **Fish** |
| Site 1  (fishponds = 139)*  (sampling-point=556)** | 19 | 120 | 47 | 92 | 44 | 164 | 60 | 188 | 112 | 484 |
| Site 2  (fishponds =143 )*  (sampling-point=572)** | 21 | 122 | 17 | 126 | 68 | 192 | 92 | 116 | 36 | 548 |
| Site 3  (fishponds =55)*  (sampling-point=220)** | 9 | 46 | 43 | 12 | 52 | 28 | 8 | 108 | 24 | 204 |

*fishpond level.

**sampling-point level.
